# The Host DHX29 RNA Helicase Regulates HCMV Immediate Early Protein Synthesis

**DOI:** 10.1101/2025.01.27.635168

**Authors:** Erik M. Lenarcic, Nathaniel J. Moorman

## Abstract

The dead box helicase DHX29 plays a critical role in the translation of mRNAs containing complex RNA secondary structure in their 5’ untranslated regions. The human cytomegalovirus (HCMV) genome has a high GC content, suggesting the 5’UTRs of viral mRNAs may contain significant secondary structure and require DHX29 for their efficient translation initiation. We found that depleting DHX29 from primary human fibroblasts prior to infection reduced viral mRNA and protein levels and decreased HCMV replication. The defect in HCMV replication correlated with decreased expression of the HCMV immediate early proteins IE1 and IE2, which are necessary for the establishment of lytic infection. Analysis of polysome associated mRNAs revealed that the defect in IE1 and IE2 expression is due to decreased mRNA translation efficiency. We found that DHX29 depletion led to reduced levels of the eIF4F translation initiation complex, resulting from decreased translation of the eIF4G mRNA. However, in line with our previous results showing a minimal role for the eIF4F complex in HCMV mRNA translation, we found that depleting eIF4G prior to infection did not impact IE1 and IE2 translation. Together our results define a new role for DHX29 in regulating eIF4F-dependent translation and identify a critical role for DHX29 in the translation of HCMV mRNAs.

**Significance:** Expression of the HCMV immediate early proteins IE1 and IE2 is critical for the establishment of lytic replication and the reactivation of latent HCMV infections. Defining the mechanisms controlling HCMV IE1 and IE2 protein expression has the potential to identify new strategies for therapeutic interventions that can limit HCMV disease in immune naïve and immune compromised individuals. Our finding that the cellular DHX29 helicase is necessary for the efficient translation of mRNAs encoding IE1 and IE2 suggests that therapies that inhibit DHX29 could potentially be useful in treating HCMV disease and adds to the growing body of literature suggesting DHX29 activity is a disease driver in multiple indications including viral disease, inflammation and cancer.

## Introduction

Human cytomegalovirus (HCMV) is a significant cause of disease in immune compromised and immune naïve individuals. Like all viruses, HCMV relies on cellular translation machinery for the expression of viral proteins. While most viruses shut off host protein synthesis to facilitate the expression of viral proteins, HCMV is somewhat unique in that cellular protein synthesis is maintained, and even increased, throughout infection(1).

The 5’ untranslated regions (5’UTR) of mRNAs play important roles in the post transcriptional regulation of protein synthesis (2). Canonical translation initiation begins with recognition of the mRNA 5’ 7-methylguanosine (m7G) cap by the eIF4E protein, which in turn recruits the eIF4G scaffold and the eIF4A DEAD box RNA helicase to form the eIF4F complex on the 5’ end of the mRNA (3). eIF4A unwinds the RNA structure in the 5’ UTR to create a landing site for the remaining members of the preinitiation complex: the 40S small ribosomal subunit, ternary complex, eIF3, eIF5, eIF1 and eIF1A. The preinitiation complex then scans the 5’ UTR until reaching the translation start codon, where the 60S large ribosomal subunit joins to form the mature 80S ribosome, and translation elongation begins. During the initiation stage, the eIF4A helicase moves along the RNA, unwinding secondary structure that would otherwise prevent the preinitiation complex from scanning the 5’UTR. However, some 5’UTRs contain extensive RNA secondary structure which is too stable for eIF4A to unwind. Such mRNAs require additional factors for their efficient translation, including DHX29 (2–4).

DHX29 interacts with the 40S ribosomal subunit and the eIF3 subunit of the preinitiation complex (5–7). On highly structured 5’ UTRs, DHX29 enhances the ability of the preinitiation complex to find the appropriate initiation codon, even when the start codon is embedded in RNA structure (7, 8). The precise mechanism by which DHX29 facilitates translation of mRNAs containing complex 5’UTRs is unknown, partly due to the discovery that this helicase is non-processive and therefore does not assist eIF4A in unwinding RNA directly (7). Evidence from experiments to determine DHX29’s influence on Internal ribosome entry site (IRES) mediated translation suggests that part of its function is to induce conformational changes in the 40S ribosomal subunit that change the way RNA secondary structure interacts with ribosomes (7).

While DHX29 plays a critical role in the translation of RNA virus genomes, its potential role in the translation of DNA virus mRNAs has not been explored. The human cytomegalovirus (HCMV) genome has a GC content of 60% (9), increasing the likelihood that the 5’UTR of any given HCMV mRNA has significant RNA secondary structure. We previously found that neither the eIF4F complex nor the eIF4A RNA helicase is required for efficient HCMV mRNA translation (10), suggesting that additional cellular and/or viral factors are needed to resolve HCMV mRNA 5’UTR structure to ensure efficient viral protein synthesis. Here we show that DHX29 is necessary for efficient HCMV lytic replication and the translation of viral mRNAs encoding the IE1 and IE2 proteins. We also show that DHX29 regulates eIF4F dependent translation by facilitating efficient eIF4G expression. However the translation of transcripts encoding IE1 and IE2 mRNA occurs is independent of eIF4G

## Methods

### Cells, plasmids and viruses

Lentivirus was produced by transfecting HEK 293T cells with either a vector encoding a scrambled control shRNA or a DHX29 specific shRNA together with MISSION Lentiviral packaging mix (Sigma) using PEI as before (10, 11). MRC5 primary human fibroblasts were transduced with lentiviruses in the presence of 8 mg/mL polybrene (hexadimethrine bromide) in Dulbecco’s Modified Eagle Medium (DMEM, Sigma) with 10% FBS incubated at 37°C and 5% CO_2_. Seventy-two hours after addition of lentivirus the cells were either mock infected or infected with the BAD*in*GFP laboratory strain of HCMV (12) in DMEM with 10% FBS incubated at 37°C and 5% CO_2_. Cell free infectious virus production during infections was quantified using a 50% tissue culture infectious dose (TCID_50_) assay as described previously (13).

### Sucrose gradient centrifugation and polysome fractionation

Analysis of polysomes was performed as described previously (10, 11, 14, 15). Briefly, cells were incubated with 100 µg/mL cycloheximide (CHX) at 37°C and 5% CO_2_ for ten minutes in DMEM and then washed twice with PBS containing 100 µg/mL CHX. The cells were then harvested by scraping into PBS plus CHX, followed by centrifugation at 1000 x g for 10 minutes. Cell pellets were resuspended in polysome gradient lysis buffer (20mM Tris-HCl, pH 7.4, 140mM KCl, 5mM MgCl_2_, 1% Triton X-100, 100 µg/mL CHX) and allowed to lyse on ice for ten minutes. The cells were then sheared by passage through a 27 gauge needle five times and nuclei were removed by centrifugation at 1100 x g for 5 minutes. The mitochondria and insoluble material were then removed by centrifugation at 21000 x g for ten minutes. The resultant post-mitochondrial supernatants were overlayed on to 10-50% linear sucrose gradients and centrifuged at 130000 x g for 2 hours. The gradient was then fractionated with continuous OD 254 monitoring using a UA-6 Absorbance Detector (Teledyne ISCO) and monitored by Peak Chart software (Brandel). To analyze proteins present in gradient fractions, a 200 µL aliquot of each gradient fraction was mixed with 1 mL of 20% trichloroacetic acid (TCA) to precipitate proteins, which were then recovered by centrifugation at 21000 x g for 30 minutes. Protein pellets were washed with acetone, resuspended in protein sample buffer (0.1 M Tris-HCl pH 6.8, 6% Glycerol, 2% SDS, 0.1 M DTT, 0.002% Bromophenol Blue), resolved by SDS-PAGE, and analyzed by Western blot.

### Quantitative reverse transcriptase PCR (qRT-PCR)

Quantification of specific RNAs was performed essentially as described previously (10, 11, 14, 15). Briefly, RNA was extracted from 200 µL aliquots of individual sucrose gradient fractions by using Trizol (Sigma) followed by Turbo DNase (Thermo Fisher) treatment. An equal volume of DNase treated RNA from each sucrose gradient fraction was used to prepare cDNA using the High Capacity cDNA Reverse Transcription Kit (Thermo Fisher) according to the manufacturer’s directions and the cDNAs were diluted 1:1 with deionized water prior to PCR. Primers for the following mRNAs were used: IE1 (CAAGTGACCGAGGATTGCAA, CACCATGTCCACTCGAACCTT), IE2 (TGACCGAGGATTGCAACGA, CGGCATGATTGACAGCCTG), β-Actin (GACCCAGATCATGTTTGAGACC, GTCACCGGAGTCCATCACGA), eIF4G (GCCATTTCAGAGCCCAACTTCTC, CGGAAGTTCACAGTCACTGTTGG), HSP90 (AGATTCCACTAACCGACGCC, CCGCACTCGCTCCACAAA) and RACK1 (TGGGATCTCACAACGGGCACCA, CCGGTTGTCAGAGGAGAAGGCCA). qPCR was performed using SYR Select Master Mix (Thermo Fisher) and 40 cycles (45 seconds at 60°C followed by 15 seconds at 95°C) in a CFX96 thermal cycler (Biorad). Total RNA was prepared and analyzed as above except that 2 ug of RNA was used to prepare cDNA instead of equal volumes.

### Protein analysis

Western blot analysis of protein expression was performed essentially as described previously (10, 11). Cells were harvested by scraping and pelleted by centrifugation at 21000 x g for 10 seconds. Cell pellets were resuspended in RIPA buffer (50 mM Tris-HCl pH 7.4, 1% NP-40, 0.25% sodium deoxycholate, 150 mM NaCl, 1mM EDTA) containing protease inhibitors (Complete EDTA-free protease inhibitor; Roche) and allowed to lyse on ice for ten minutes. Insoluble material was removed by centrifugation at 21000 x g for ten minutes at 4°C and the protein concentration in the supernatant was determined by the Bradford assay. 30 µg per sample were resolved by SDS-PAGE gel electrophoresis and transferred to nitrocellulose membranes. Membranes were probed with antibodies to DHX29 (Santa Cruz Biotechnology), eIF4E (Cell Signaling Technology), β-Actin (Santa Cruz Biotechnology), rpS6 (Cell Signaling Technology), rpL13a (Cell Signaling Technology), HCMV IE1, IE2, UL44, UL99 (pp28), eIF4G (Cell Signaling Technology), 4E-BP (Cell Signaling Technology), 4E-BP-phospho (Cell Signaling Technology), rpS6-phospho (Cell Signaling Technology) and HSP90 (Stressgen).

### Cap binding complex precipitation

Isolation of m7G associated proteins was performed essentially as described previously (10, 16). Cells pellets were resuspended in CAP IP buffer (40mM HEPES, pH 7.6, 120mM NaCl, 1mM EDTA, 0.3% CHAPS, Complete EDTA-free protease inhibitor) and allowed to lyse on ice for 10 min then insoluble material was removed by centrifugation at 21000 x g for ten minutes at 4°C. The protein concentration of each supernatant was analyzed by Bradford and equal amounts of protein per condition were used in each experiment. Some samples were incubated with 100 µM soluble 7-methyguanosine 5’-triphosphate (Sigma) for one hour at 4°C with nutation. Samples were then mixed with immobilized γ-Aminophenyl-m^7^GTP agarose breads (Jena Bioscience) and incubated with nutation for one hour at 4°C. The beads were pelleted by centrifugation at 6000 x g for one second and washed with CAP IP buffer three times. The beads were then resuspended in protein sample buffer and boiled at 100°C for ten minutes. The supernatants were then separated by SDS-PAGE and analyzed by Western blot.

### Metabolic ^35^S-labeling and immunoprecipitation

Cells were incubated in growth media lacking methionine and cysteine (Sigma) for fifteen minutes at 37°C and 5% CO_2_, then 125 µCi of EasyTag Express^35^S Protein Labeling Mix (Revvity) was added per mL of growth media and the cells were incubated at 37°C and 5% CO_2_ for an additional thirty minutes. The cells were then washed twice with PBS and harvested, followed by lysis in RIPA buffer. Lysates were pre-cleared prior to immunoprecipitation by incubation for thirty minutes at 4°C with Protein A/G PLUS-Agarose beads (Santa Cruz Biotechnology) with constant mixing. Then HCMV IE1 protein was precipitated from pre-cleared lysates by incubation for one hour at 4°C with IE1 antibody pre-conjugated to protein A/G beads with constant mixing. The pre-conjugated beads were then washed three times with RIPA buffer before being resuspended in sample buffer and boiled at 100°C for ten minutes. Proteins eluted from the beads were resolved by SDS-PAGE and analyzed by autoradiography (GeneMate).

## Results

### HCMV infection increases the association of DHX29 with translation machinery during HCMV infection

Viruses, including HCMV, manipulate initiation factors during lytic infection to enhance viral mRNA translation (17–23). The RNA helicase DHX29 participates in translation initiation in uninfected cells (24), however its role in translation during HCMV infection was unknown. To determine if DHX29 associates with proteins involved in translation initiation during infection, we measured the association of DHX29 with 7-methylguanosine (m^7^G) sepharose in extracts of HCMV infected cells. Similar levels of the eIF4E, the 5’ m^7^G cap binding protein precipitated with m^7^G beads incubated with lysates of uninfected and infected cells (Fig. 1A). However, HCMV infection increased the amount of DHX29 associated with m^7^G beads despite similar overall levels of DHX29 in uninfected and infected cell lysates. DHX29 association with m^7^G beads was specific, as the inclusion of excess soluble m^7^G blocked the co-purification of both eIF4E and DHX29 with the m^7^G beads. These results suggest that HCMV infection increases the association of DHX29 with the 5’ mRNA cap.

**Figure 1:**
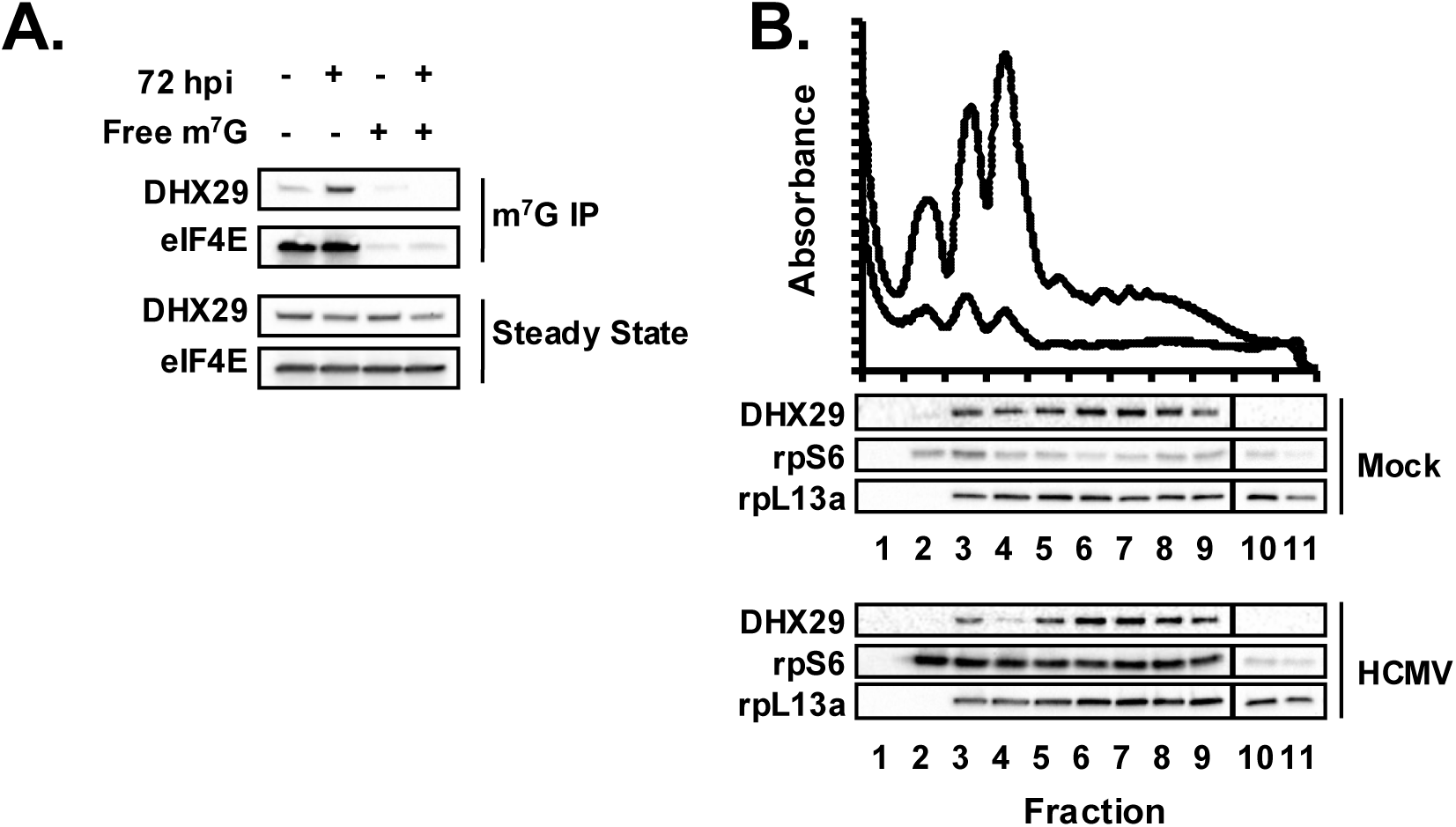
DHX29 associates with the mRNA cap and actively translating mRNAs during HCMV infection. A) Proteins co-purifying with m^7^G sepharose were analyzed by Western blot at 72 hours after HCMV infection (MOI of 3). Excess free m^7^G was included in some reactions (free m^7^G) to control for specificity. B) Cytoplasmic lysates from uninfected (Mock, solid line) or HCMV infected (72hpi, MOI of 3, dashed line) fibroblasts were resolved through sucrose density gradients to separate ribosomal subunits (40S, 60S), single ribosomes (80S), and polysomes. Proteins extracted from each gradient fraction were analyzed by Western blot. The results are representative of at least three independent experiments.

We also measured the association of DHX29 with actively translating mRNAs during infection by measuring DHX29 association with polysomes in sucrose density gradients. As before, we found that HCMV infection increases the abundance of polysomes as compared to uninfected cells (Fig. 1B), consistent with the increase in overall levels of protein synthesis caused by HCMV infection (17, 18, 25–30). DHX29 was detected in fractions containing ribosomal subunits, monosomes and polysomes in both uninfected and infected cytoplasmic extracts. However, DHX29 was more concentrated in fractions containing polysomes in extracts from infected cells. Together with the increased association of DHX29 with the m^7^G mRNA cap, these results suggest an enhanced role for DHX29 in mRNA translation during HCMV infection.

### DHX29 is required for efficient HCMV amplification

The increased association of DHX29 with translation initiation complexes and translating mRNAs during infection suggested it may play an important role in HCMV replication. To determine the role of DHX29 in HCMV replication, we depleted DHX29 from human fibroblasts and then measured the effect on the production of infectious HCMV virions. DHX29 depletion decreased the amount of infectious HCMV produced by >10 fold (Fig. 2A). The defect in virus replication correlated with delayed and reduced expression of viral immediate early (IE1, IE2), early (UL44), and late (pp28) proteins throughout the replication cycle (Fig. 2B). Together these data show that DHX29 is critical for efficient viral protein expression and, as a result, HCMV replication.

**Figure 2:**
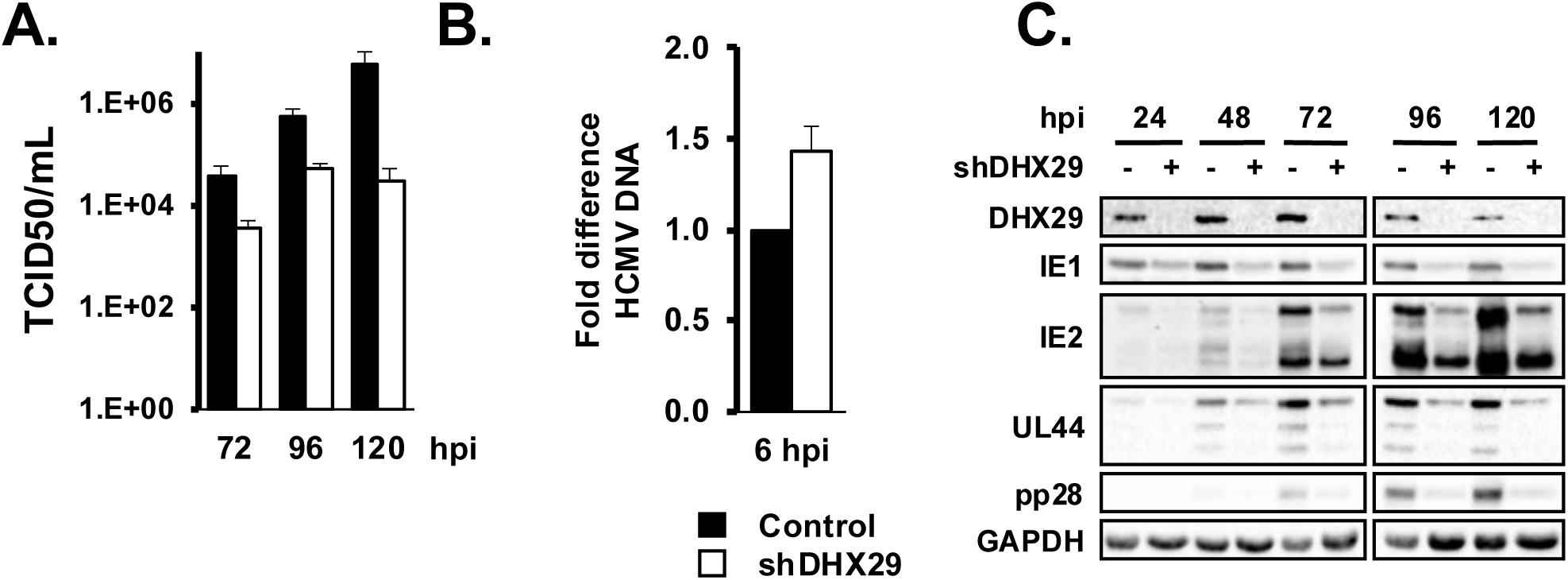
DHX29 is necessary for efficient HCMV replication. A) Human fibroblasts were transduced with lentivirus encoding scrambled (control) or DHX29 specific shRNAs. Seventy -two hours later cells were infected with HCMV (MOI of 3). Cell free virus was harvested at the indicated times after infection and quantified by the TCID50 assay. B) Cells were transduced and infected as in A. The amount of HCMV DNA in infected cells was measured at six hours post infection by qPCR. The mean data from three independent experiments is shown. C) Cells were transduced and infected as in A, and viral protein expression was measured by Western blot at the indicated times after infection. The data are representative of at least three independent experiments.

### DHX29 is necessary for the efficient translation of HCMV immediate early mRNAs

The decrease in immediate early protein expression suggested that DHX29 is required for a very early step in the HCMV infectious cycle. DHX29 was not required for efficient HCMV entry, as similar numbers of viral genomes were found in infected cells at 6 hours after infection of control or DHX29-depleted fibroblasts (Fig. 2B). Similarly, viral immediate early mRNA levels were not significantly affected by DHX29 depletion (Fig. 3A). In contrast, the expression of the HCMV immediate early proteins IE1 and IE2 was decreased in the absence of DHX29 (Fig. 3B).

**Figure 3.**
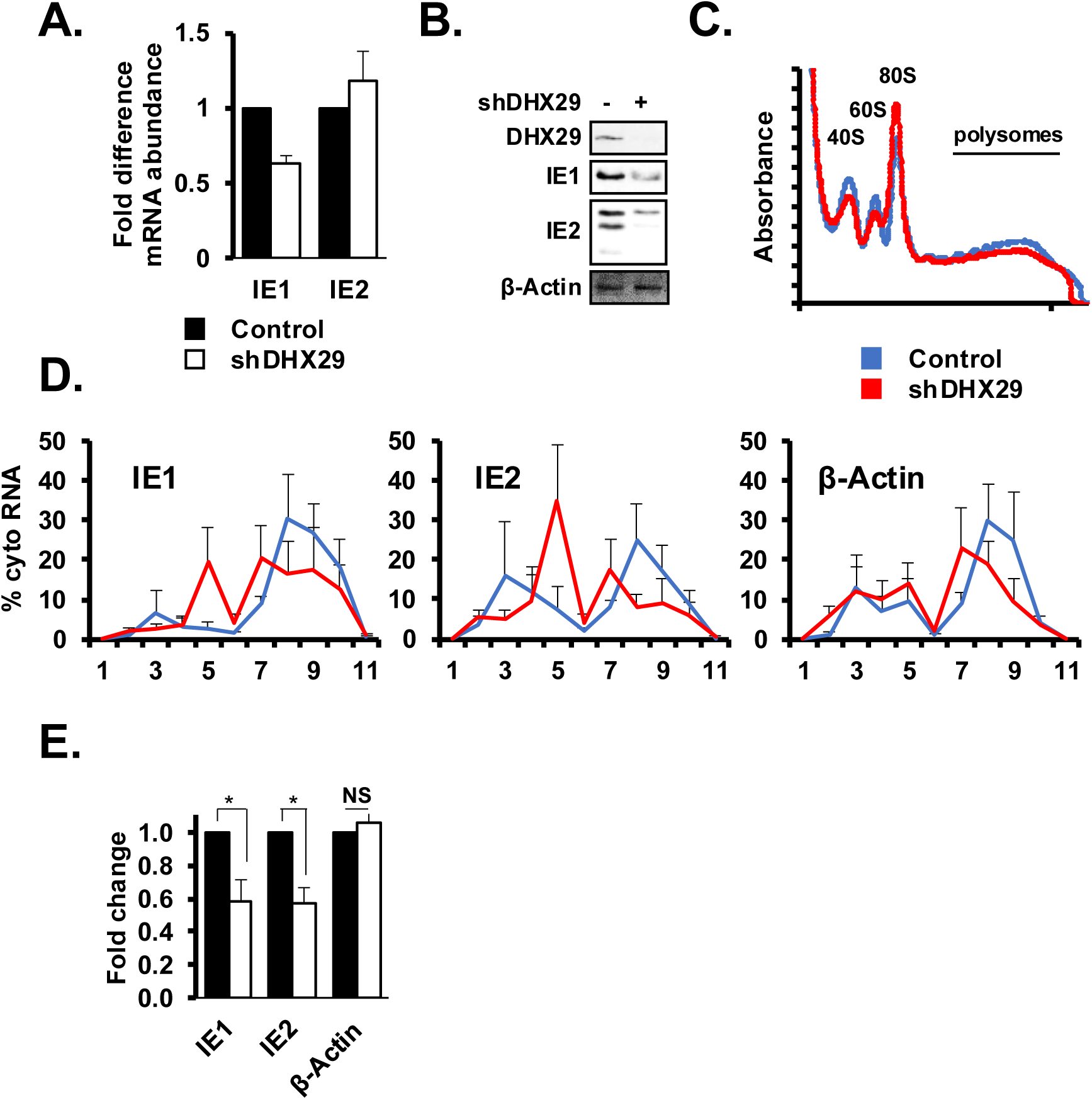
DHX29 is necessary for the efficient translation of mRNAs encoding HCMV immediate early proteins. Human fibroblasts transduced with lentivirus encoding scrambled control or DHX29 specific shRNAs were infected with HCMV (MOI of 3). Six hours post infection HCMV immediate early mRNA (A) and protein levels (B) were analyzed by qRT-PCR and Western blot, respectively (B). C) Polysome formation in control or DHX29-depleted cells was measured by sucrose gradient centrifugation at 6 hours after HCMV infection. D) The percent of cytosolic RNA found in each gradient fraction was measured by qRT-PCR for the IE1, IE2, and β-actin mRNA. E) Graph of data in D, comparing the amount of each mRNA found in gradient fractions containing polysomes in control or DHX29 -depleted cells. The amount of each mRNA in control cells was set to 100%. The mean data from 3 independent experiments is shown. (* p<0.05; ** p<0.005; *** p<0.001)

The decrease in HCMV IE protein expression despite similar mRNA levels suggested a role for DHX29 in the translation of the HCMV IE mRNAs. To determine if DHX29 was necessary for efficient HCMV IE mRNA translation, we measured the amount of IE1 and IE2 mRNA associated with polysomes in control or DHX29-depleted fibroblasts. Cytoplasmic lysates from HCMV infected cells were resolved through linear sucrose gradients to measure the abundance of ribosomal subunits, ribosomes, and polysomes in the presence of DHX29. Depleting DHX29 did not affect the overall abundance of polysomes at 6 hours after infection, indicating that DHX29 depletion did not affect overall levels of translation (Fig. 3C).

To determine the effect of DHX29 depletion on HCMV IE mRNA translation, we quantified the amount of the IE1 and IE2 mRNAs associated with polysomes in control or DHX29-depleted cells. Both the IE1 and IE2 mRNAs were less abundant in gradient fractions containing polysomes, and more abundant in fractions containing monosomes, when DHX29 levels were reduced (Fig. 3D). The amount of IE1 and IE2 mRNAs associated with polysomes was consistently reduced by 50% in DHX29 depleted cells (Fig. 3E). In contrast, DHX29 depletion did not affect the distribution of the host β-actin mRNA in the gradient, and the amount of β-actin mRNA associated with polysomes was unchanged (Fig. 3D, E). This data shows that DHX29 is required for the efficient translation of HCMV IE1 and IE2 mRNAs.

### DHX29 depletion inhibits eIF4G dependent translation

For some mRNAs DHX29 enhances translation in cooperation with the eIF4F translation initiation complex (7). We therefore tested the effect of DHX29 depletion on eIF4F complex formation. Proteins associated with the m^7^G mRNA cap were captured from infected cell extracts from control or DHX29-depleted cells using m^7^G sepharose and the presence of eIF4F subunits was analyzed by Western blot. While eIF4E was recovered at similar levels from control and DHX29 depleted cells, the amount of the eIF4G protein associated with the beads was reduced in the absence of DHX29 (Fig. 4A). The decrease in eIF4G binding was not due to decreased mTOR activity in the absence of DHX29, as the levels of phosphorylated 4E-BP and rpS6 were similar to those in infected control fibroblasts (Fig. 4D). Rather, the decrease in eIF4G association with m^7^G sepharose correlated with reduced steady state levels of eIF4G in DHX29-depleted cells (Fig. 4A).

**Figure 4.**
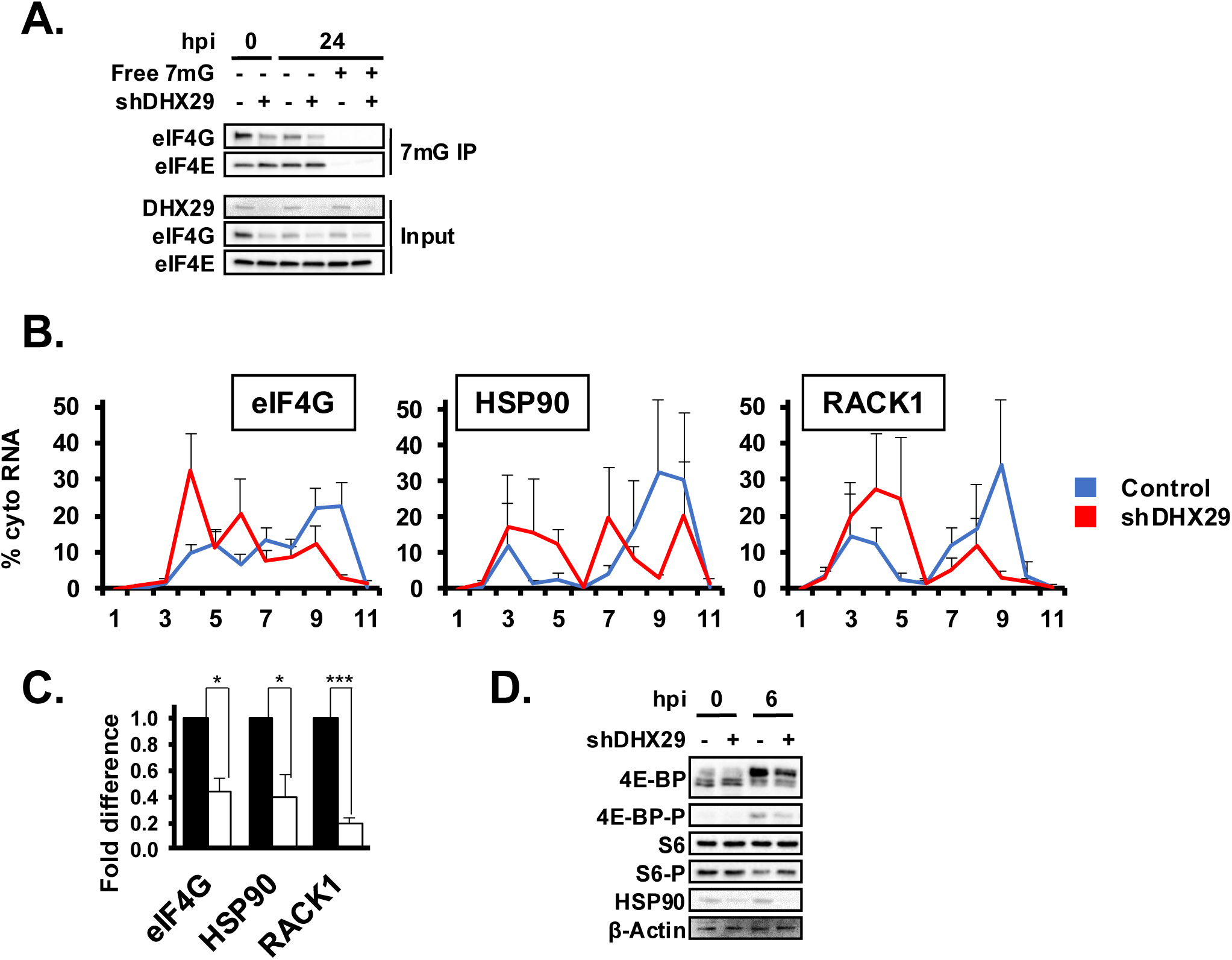
DHX29 depletion disrupts eIF4F by reducing eIF4G expression without affecting mTOR activity. A) Human fibroblasts transduced with lentivirus expressing scrambled control or DHX29 specific shRNAs were infected with HCMV (MOI of 3). Six hours post infection m^7^G cap binding complexes were precipitated from cell lysates and analyzed by Western blot. B) Cells were treated as in A, and cytoplasmic extracts were fractionated by sucrose gradient centrifugation. The percent of the eIF4G, HSP90, or Actin in each gradient fraction was measured by qRT-PCR. C) Graph comparing the amount of each RNA in gradient fractions containing polysomes in control or shDHX29 expressing cells. The amount of each mRNA in control cells was set to 100%. D) Western blot showing the phoshorylation state of the mTOR substrates 4E-BP1 and rpS6 in control or DHX29 depleted cells. The mean data from 3 independent experiments is shown. (* p<0.05; ** p<0.005; *** p<0.001)

The eIF4G mRNA has a complex 5’ UTR (31, 32), suggesting that the defect in eIF4G protein expression in the absence of DHX29 may be due to decreased translation. Using sucrose density centrifugation, we found that DHX29 depletion resulted in an increase in the amount of eIF4G mRNA migrating in fractions of the gradient containing monosomes and ribosomal subunits, with a concomitant 50% reduction in the amount of eIF4G mRNA in gradient fractions containing polysomes (Fig. 4B). If DHX29 is necessary for eIF4G expression, then eIF4F dependent mRNA translation should be reduced when DHX29 is depleted. Consistent with this hypothesis, DHX29 depletion reduced the association of the eIF4F-dependent HSP90 and RACK1 mRNAs with polysomes (Fig. 4C and ref(10)), resulting in a 50% and 60% reduction in their translation efficiency, respectively (Fig. 4D). These data show that efficient eIF4G mRNA translation, and thus eIF4F complex formation, requires DHX29 expression. Further, these data suggest that DHX29 is necessary for the efficient translation of eIF4F-dependent mRNAs.

### Reduced HCMV immediate early protein synthesis is not due to reduced eIF4G expression

We previously found that inhibition or disruption of eIF4F did not impact IE1 translation (10), suggesting that a reduction in eIF4G expression and eIF4F complex formation when DHX29 is depleted cannot explain the defect in IE1 and IE2 protein expression. However, eIF4G can also facilitate translation independent of the eIF4F complex on some mRNAs (22, 23). To determine if reduced eIF4G expression could explain the defect in IE1 and IE2 translation in the absence of DHX29, we measured the effect of eIF4G depletion on IE1 and IE2 protein levels. In addition, we treated eIF4G-depleted cells with the mTOR inhibitor Torin1 (Fig. 5A), which activates the 4E-BP translation repressor and blocks eIF4G association into the eIF4F complex (10, 13, 33). Torin1 treatment of eIF4G-depleted fibroblasts enhanced 4E-BP binding to the m^7^G sepharose as expected. However, IE1 and IE2 protein levels were unaffected by eIF4G depletion alone or with Torin1 treatment. As an additional measure of the role of eIF4G in HCMV IE1 and IE2 expression, we measured IE1 and IE2 protein levels and rates of synthesis in cells where eIF4G had been inactivated by prior infection with human rhinovirus (Fig. 5B), which encodes a protease that cleaves eIF4G thereby disrupting its interaction with eIF4E. The steady state protein levels of IE1 and IE2 were unaffected by prior human rhinovirus (HRV) infection despite efficient eIF4G cleavage. In addition, the same amount of IE1 and IE2 was synthesized during the final thirty minutes of the experiment in the presence or absence of HRV infection (Fig. 5C). These data demonstrate that eIF4G, like the eIF4F complex itself, is not necessary for the efficient translation of the IE1 and IE2 mRNAs. Further, these data suggest that the defect in IE1 and IE2 mRNA translation in the absence of DHX29 is not the result of decreased eIF4G expression.

**Figure 5.**
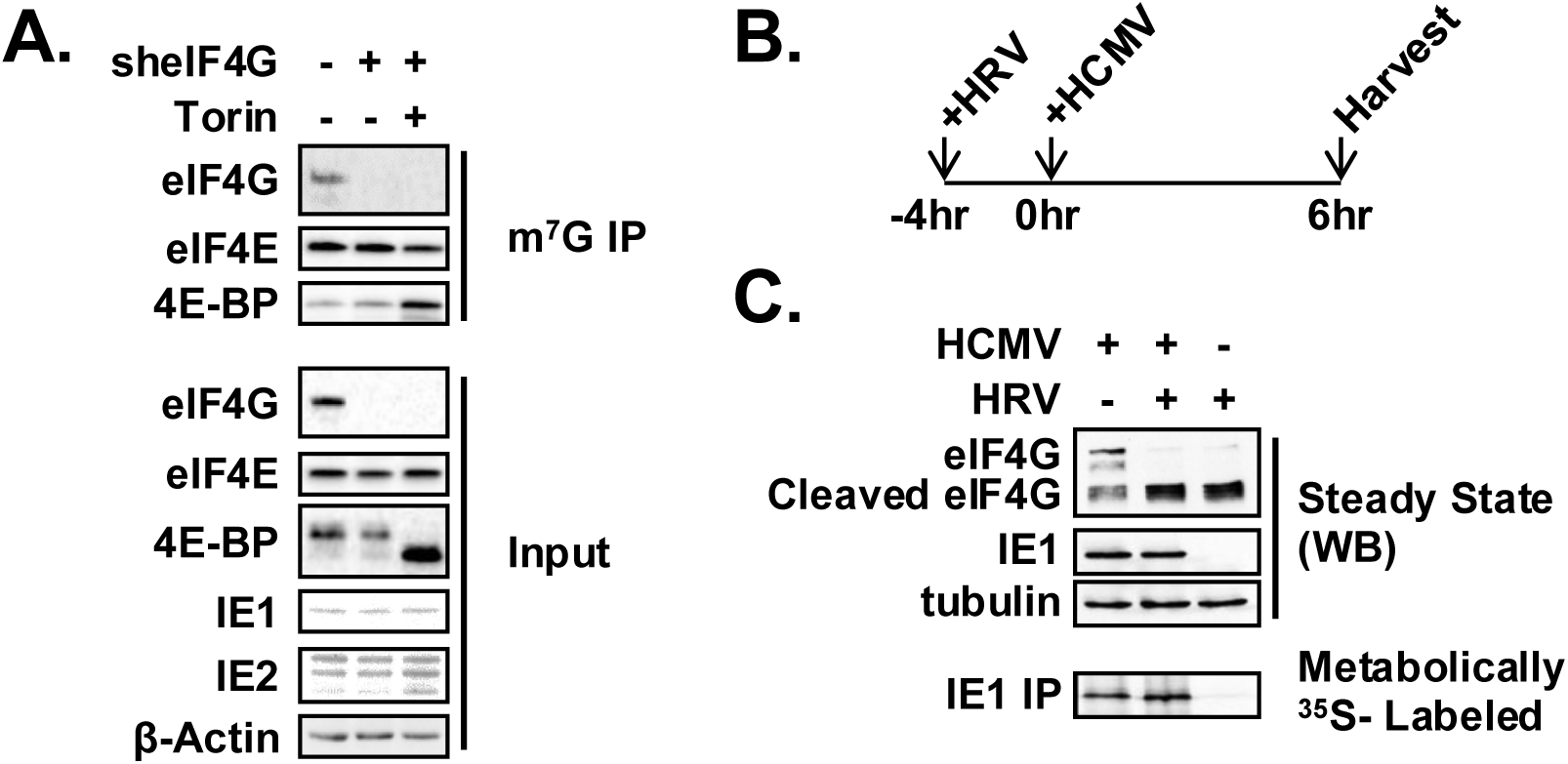
Efficient HCMV IE protein expression does not require eIF4G. A) Human fibroblasts transduced with lentivirus encoding scrambled control or eIF4G specific shRNAs were infected with HCMV. Six hours post infection cap binding complexes were precipitated from cell lysates (m^7^G IP) and analyzed by western blot along with input lysates (steady state). B) Cartoon explaining the effect of Human Rhinovirus (HRV) 2A protease on translation initiation. C) Human fibroblasts were infected with HRV, followed by HCMV infection four hours later (top). Six hours post infection proteins were analyzed by Western blot or were metabolically labeled with ^35^S-methionine for thirty minutes before being immunoprecipitated with an anti-IE1 antibody and analyzed by gel electrophoresis and autoradiography (bottom).

## Discussion

The role of DHX29 in translation initiation continues to be studied and refined. Here we show that DHX29 plays an important role in HCMV infection. We found that DHX29 is required for efficient HCMV replication and associates with the translation machinery in HCMV infected cells. In the absence of DHX29, the IE1 and IE2 mRNAs are translated less efficiently. In addition, DHX29 depletion decreases the expression of the eIF4G component of the eIF4F translation initiation complex, which is required for the expression of cellular proteins necessary for efficient HCMV replication (25). However, the defect in IE1 and IE2 expression was independent of the effect on eIF4G, as both IE1 and IE2 proteins were efficiently expressed when eIF4G was depleted or inactivated, consistent with previous studies showing a minimal role for the eIF4F translation initiation complex in the translation of HCMV mRNAs. Together these results suggest that DHX29 has two roles in HCMV replication – directly supporting the translation of viral mRNAs as well as the expression of cellular proteins needed for virus replication.

These data raise the question of why DHX29 is needed for translation of the IE1 and IE2 mRNAs. DHX29 is required for the efficient translation of mRNAs containing moderate to extensive structure (ΔG < -40kcal/mol) in their 5’UTRs (24). Transcripts encoding IE1 and IE2 that originate from the major immediate early promoter (MIEP) contain a 136 nt 5’UTR with a predicted free energy of < -40 kcal. Similarly, the 5’UTR of transcripts encoding eIF4G1 are 368 nt in length (31, 32) and are predicted to contain highly structured regions with free energies around -100 kcal/mol (31, 34), potentially explaining the requirement for DHX29 for efficient eIF4G1 protein expression. In contrast, the actin mRNA 5’UTR has a predicted free energy of < -16 kcal/mol and does not require DHX29 for its translation (Figure 3 and ref(24)). The nature of the 5’UTR of the IE1 and IE2 mRNAs therefore likely explains the need for DHX29 for their translation and likely explains the role for DHX29 in eIF4G expression. While the full-length mRNA structure for most HCMV transcripts remains to be defined, the high GC content of the HCMV genome suggests DHX29 may be similarly required for the efficient translation of additional viral mRNAs as well.

Our results also raise the question of how DHX29 facilitates the translation of transcripts encoding IE1 and IE2. DHX29 is a non-processive RNA helicase whose activity is enhanced by binding to the 40S ribosomal subunit (7). DHX29 is located at the mRNA entry channel of the 48S translation initiation complex where it interacts with helix 16 of the 18S rRNA and specific components of the eIF3 complex (6, 35). From its position in the 48S complex DHX29 is thought to resolve RNA structures that otherwise impede mRNA transit through the entry channel of the ribosome (36). Interestingly the 5’UTR of mRNAs encoding IE1 and IE2 contain a stable hairpin immediately 5’ of the AUG initiation codon (14). One potential model explaining the role of DHX29 in IE1 and IE2 expression is that DHX29 associates with 48S complexes bound to mRNAs encoding IE1 and IE2 and enhances unwinding of the AUG-proximal hairpin to facilitate 48S scanning of the 5’UTR and ensure correct recognition of the IE1 and IE2 start codon.

Our results also shed further light on the role of the eIF4F initiation complex in the translation of HCMV mRNAs. For many mRNAs, translation begins with the nucleation of the tripartite eIF4F complex on the mRNA 5’ cap. The eIF4E protein binds the cap and recruits the eIF4G scaffold protein, which recruits the 43S preinitiation complex, and the eIF4A RNA helicase, which unwinds RNA structures to enhance scanning (37). Our group and others previously found that viral mRNA translation is largely resistant to eIF4F disruption and depletion or inhibition of eIF4A (10, 11, 38). Here we find that eIF4G abundance and functionality are also dispensable for IE1 and IE2 expression. IE1 and IE2 protein synthesis was unaffected by inactivation of full-length eIF4G by infection with human rhinovirus, which encodes a protease that cleaves eIF4G. In specific instances an eIF4G cleavage product can functionally substitute for full-length eIF4G. However, that was likely not the case here as depletion of eIF4G prior to infection was similarly ineffective at reducing IE1 and IE2 expression, even when residual eIF4F complex was disrupted by treatment with the mTOR inhibitor Torin1. The reduction in eIF4G expression is therefore not the cause of the reduction in IE1 and IE2 expression, however its effect on viral amplification may not be inconsequential, as eIF4F plays a critical role in the translation of cellular mRNAs that HCMV requires for replication (25). These data provide further support for a non-canonical translation initiation process on HCMV mRNAs independent of eIF4F or its components.

Interestingly, DHX29 regulates translation initiation for other viruses as well. For example, Sindbis virus 26S RNA required DHX29 for efficient initiation complex formation (39). Other viruses use alternatives to cap dependent translation initiation that may also be influenced by DHX29. Small positive stranded RNA viruses typically have structured 5’ UTRs, many of which initiate translation cap independently using an IRES. A subset of picornavirus IRESes have been shown to require DHX29 for activity (8, 40). We have described the first requirement for DHX29 on a transcript from a DNA virus. However, the breadth of viral transcripts requiring DHX29 during HCMV infection cannot be determined by shRNA knockdown since defects arise so early after infection. Further analysis by other approaches will be needed to elucidate the extent to which HCMV protein expression relies upon DHX29.

## Acknowledgements

The authors wish to acknowledge members of the Moorman lab for helpful conversations and discussions. This work was supported by grants R01-AI103311 and R21 R21-AI123811 to N.J.M from the National Institute of Health, and the University of North Carolina Cancer Research Fund.

## Notes

### Competing Interest Statement

The authors have declared no competing interest.

